# Encapsulation and recovery of murine hematopoietic stem and progenitor cells in a thiol-crosslinked maleimide-functionalized gelatin hydrogel

**DOI:** 10.1101/2021.03.22.436491

**Authors:** Aidan E. Gilchrist, Julio F. Serrano, Mai T. Ngo, Zona Hrnjak, Sanha Kim, Brendan A.C. Harley

**Author notes:** To whom correspondence should be addressed: B.A.C. Harley, Department of Chemical and Biomolecular Engineering, Cancer Center at Illinois, Carl R. Woese Institute for Genomic Biology, University of Illinois at Urbana-Champaign, 110 Roger Adams Laboratory, 600 S. Mathews Ave., Urbana, IL 61801.

## Abstract

Biomaterial platforms are an integral part of stem cell biomanufacturing protocols. The collective biophysical, biochemical, and cellular cues of the stem cell niche microenvironment play an important role in regulating stem cell fate decisions. Three-dimensional (3D) culture of stem cells within biomaterials provides a route to present biophysical and biochemical stimuli such as cell-matrix interactions and cell-cell interactions via secreted biomolecules. Herein, we describe a maleimide-functionalized gelatin (GelMAL) hydrogel that can be crosslinked via thiol-Michael addition click reaction for the encapsulation of sensitive stem cell populations. The maleimide functional units along the gelatin backbone enables gelation via the addition of a dithiol crosslinker without requiring external stimuli (*e.g*., UV light, photoinitiator), reducing reactive oxide species generation. Additionally, the versatility of crosslinker selection enables easy insertion of thiol-containing bioactive or bioinert motifs. Hematopoietic stem cells (HSCs) and mesenchymal stem cells (MSCs) were encapsulated in GelMAL, with mechanical properties tuned to mimic the *in vivo* bone marrow niche. We report insertion of a cleavable peptide crosslinker that can be degraded by the proteolytic action of SortaseA, a mammalian-inert enzyme. Notably, SortaseA exposure preserves stem cell surface markers, an essential metric of hematopoietic activity used in immunophenotyping. This novel GelMAL system enables a route to producing artificial stem cell niches with tunable biophysical properties with intrinsic cell-interaction motifs and orthogonal addition of bioactive crosslinks.

## 1. Introduction

Stem cells are defined by their capacity to produce multiple different cell types and self-renew [1]. Stem cells are essential for maintaining multicellular systems and, therefore, are vital for clinical therapies to regenerate or repair entire tissues [2-5]. Mesenchymal stem cells (MSCs), for example, can regenerate multiple types of tissue, are immunomodulatory, and have been extensively explored for regenerative therapies, with over 950 clinical trials currently registered with the FDA [6-8]. Hematopoietic stem cell (HSC) transplants are perhaps the most widely used stem cell regenerative therapy and are used to treat disorders of the blood and immune systems, with over 24,000 HSC transplants in 2018 in the U.S. [9-12]. These examples showcase the potential of stem cells for regenerative medicine; however, the utility of stem cell therapies has been undercut by a requirement for large cell numbers coupled with the rarity of stem cells [13, 14]. Consequently, there is a need for *in vitro* culture systems for stem cells in order to condition or expand these rare populations prior to utilization [13-15].

Biomaterials offer an advantageous route towards stem cell culture by providing a biophysical and biochemical structure that can potentially mimic the *in vivo* stem cell microenvironment or niche. Encapsulation of cells within a 3D hydrogel enables incorporation of extracellular matrix-inspired ligands, mechanotransducive stimuli, and delivery of soluble factors [16-18]. Additionally, soluble factor and matrix interactions offer an intrinsic means to control cell-cell communication by mediating passive or active transport of cell-secreted factors [19-22]. In our lab we have demonstrated engineered domains of autocrine and paracrine signaling in hydrogels to mediate HSC behavior [23-25]. Using hydrogels with restricted or enhanced cell-cell communication (autocrine vs. paracrine signaling) we, and others, have identified soluble factors such as TGFβ-1 and SCF implicated in stem cell fate decisions [26-29]. In addition, by presenting covalently-bound SCF or the cell adhesive motif, RGD, hydrogels led to increased HSC maintenance [30, 31]. Culture in 3D hydrogel also offers additional tunable variables such as nano/macrostructure, chemistry, and biophysical properties [32, 33].

Synthetic polymers, notably polyethylene-glycol (PEG), have been widely used for stem cell culture, as it provides a bioinert substrate with orthogonal control of material parameters, *e.g*., mesh size, mechanics, and bioactivity. However, synthetic polymers typically require additional functionalization through peptide linkers or interaction sites for cell viability and cell-matrix interactions [34-37]. Conversely, biopolymers such as fibronectin and gelatin, which are derived from natural materials, offer inherent cell interaction motifs. Gelatin is a commonly used biopolymer for stem cell culture [38-40] as it possesses intrinsic cell interaction motifs, including cell-adhesive RGD sites and degradation motifs for material remodeling [41, 42], which have been linked to enhanced stem cell maintenance and expansion [31, 43-45]. A major challenge facing gelatin hydrogel fabrication is the thermally reversible physical gelation process, which requires a low temperature (<30°C) to maintain a gel [46-48]. To circumvent this, the gelatin peptide backbone can be functionalized with chemical crosslinking sites, allowing for a chemically-crosslinked hydrogel that is stable at biologically relevant temperatures (37°C) [40, 49, 50]. Methacrylamide-functionalized gelatin (GelMA) has become widely used [38, 41, 51-55], with hydrogel networks formed via covalent radical-initiated crosslinking of pendulant methacryloyl groups functionalized along the gelatin backbone. However, this process is a potential drawback to cell culture as it requires UV light and a free-radical photoinitiator. Although small doses of UV exposure have been shown not to affect viability of cell lines, there is concern that the initial burst of radical formation during photopolymerization will produce reactive oxygen species (ROS) [56]. The sensitivity of primary stem cells and the role of ROS in stem cell fate decisions [57] highlights the need for a hydrogel system without the need for UV or radical crosslinking. Additionally, although chemical crosslinks are an effective means of generating a crosslinked hydrogel network, they can hinder cell recovery, as enzymatic methods are required to cleave the gelatin backbone [58]. These methods negatively impact cell viability and can cause internalization of sensitive surface markers which are essential for downstream phenotyping of stem cell populations [59, 60].

Beneficial modifications to a gelatin biomaterial system would enable encapsulation of stem cells with intrinsic material-cell interactions without the introduction of confounding ROS. Such a platform should also allow for the safe and gentle recovery of cultured stem cells without disruption of markers required for downstream analysis. Herein, we use HSCs and MSCs to demonstrate a novel gelatin system that leverages inherent cell-interaction motifs, high stem cell viability, and full recovery of stem cell markers. We describe an approach to generate maleimide-functionalized gelatin (GelMAL) hydrogels with tunable mechanics and the capacity to incorporate peptides through thiol-maleimide Michael addition click reaction without the generation of free radicals. We also report insertion of degradable dithiol peptides for complete recovery of cell surface markers. HSCs are a model system that demonstrates the utility of GelMAL, as ROS play an important role in hematopoiesis by influencing HSC self-renewal, differentiation, and cell death [61-64]. Furthermore, robust preservation of surface markers is essential for immunophenotyping of hematopoietic lineage cells [65, 66].

## 2. Methods and Materials

### 2.1 Synthesis of GelMAL Biopolymer

In a 20-mL glass vial, 0.25 g of Type A porcine gelatin (Sigma-Aldrich Company, Saint Louis, MO) was dissolved in a mixture of 5:4 (mL:mL) DI water:DMSO (Sigma-Aldrich, Saint Louis, MO) at 40 °C. Then, 3-(maleimido)propionic acid *N*-hydroxysuccinimide ester (prepared as previously reported [67]) (0.125 g, 0.47 mmol) was dissolved in 1 mL of DMSO and added to the dissolved gelatin. After stirring at 40 °C for 24 h, the resulting light pink solution was diluted with 10 mL of DI water and purified using dialysis membrane (MW cutoff 3.5 kDa, Spectrum Laboratories, Rancho Dominguez, CA) in DI water at 40 °C for 5 d. Samples were aliquoted into 5-mL batches, lyophilized, and stored at −20 °C until further use.

### 2.2 NMR Spectroscopy of Gelatin and GelMAL

#### 2.2.1 ^1^H HMR Spectroscopy

Degree of functionalization of gelatin and GelMAL was calculated using ^1^ H NMR spectrometry. Samples were prepared by dissolving approximately 15 mg of biopolymer in 700 µL of deuterium oxide (Sigma Aldrich, St. Louis, MO). NMR spectra were measured at ambient temperature on a Bruker Carver B500 spectrometer (500 MHz). Prior to the analysis and interpretation, phase correction was applied to all spectra to obtain purely absorptive peaks, baseline correction was applied before integrating the signals of interest, and the chemical shift scale was adjusted by referencing ^1^H NMR chemical shifts to the residual solvent peak at 4.80 ppm in D_2_O. GelMAL ^1^ H NMR (500 MHz, D_2_O,): *δ* 7.25 (m), 6.80 (s). Degree of functionalization (DOF) was defined as the number of modified lysine groups. Briefly, the aromatic integral was set to 1 in each spectrum and the peak area of lysine methylene protons was used for calculation of the DOF(%) as *DOF = (1 - the area of lysine methylene of GelMAL/ the area of lysine methylene of gelatin)x100*.

#### 2.2.2 ^1^H−^13^C-HSQC (Heteronuclear Single Quantum Coherence) NMR Spectroscopy

Gelatin and GelMAL samples for 2D NMR spectroscopy were prepared by dissolving approximately 15 mg of biopolymer in 700 μL of deuterium oxide. High-resolution 500 MHz NMR spectra were measured at ambient temperature on a Bruker Carver B500 spectrometer (500 MHz). Water suppression was not applied to prevent the distortion of nearby peaks. Prior to the analysis and interpretation, phase correction was applied to all spectra to obtain purely absorptive peaks, baseline correction was applied before integrating the signals of interest, and the chemical shift scale was adjusted by referencing ^1^H NMR chemical shifts to the residual solvent peak at 4.80 ppm in D_2_O.

### 2.3 Hydrogel formation

#### 2.3.1 DTT hydrogels

A 3 or 5 wt% solution of GelMAL was prepared by dissolving GelMAL in PBS (Gibco Invitrogen, Thermo Fisher Scientific, Waltham, MA) and magnetically stirring the solution in a glass vial at 45 °C, yielding a very light pink solution. Hydrogels were prepared by dispensing 100 µL of the GelMAL solution into a Teflon mold (10 mm diameter) and rested at 4 °C for 25 min to allow for physical gelation. Then, the hydrogels were removed from the molds and placed into wells containing freshly prepared 1.25 mg/mL aqueous solution of 1,4-dithiothreitol (DTT, Sigma-Aldrich, Saint Louis, MO) on a shaker at RT for 8 min. The DTT solution was removed from the wells, and the hydrogels were washed in PBS (1 mL) for 5 min (PBS wash repeated two more times). For MSC viability and behavior studies, 3 wt% and 5 wt% GelMAL hydrogels were prepared as above, with hMSCs resuspended at a density of 10^6^ MSC/mL in GelMAL precursor solution prior to dispensing into a Teflon mold (5 mm diameter, 20 µL) and crosslinking with DTT.

#### 2.3.2 Peptide and PEG dithiol hydrogels

For peptide-crosslinked hydrogels, a 10 mM solution of a custom-made peptide (GCRD-LPRTG-DRCG, Thermo Fisher Scientific, Waltham, MA) was prepared in PBS degassed with argon for 3 h. The LPRTG motif is an uncommon sequence in mammalian proteins and is specifically recognized and cleaved by a bacterial enzyme, SortaseA [68]. For PEG-dithiol crosslinked hydrogels, a 10 mM solution of PEG-dithiol (M_n_ = 1,000, Sigma Aldrich) was prepared in PBS. Hydrogels for mechanical testing were prepared by dispensing 75 µL of a concentrated GelMAL solution (6.66 wt%) into a Teflon mold (10 mm diameter, 100 µL). Crosslinking was initiated by addition of 25 µL peptide or PEG-dithiol solution for a final concentration of 5 wt% GelMAL and 2.5 mM crosslinker. The filled molds were kept at RT (25 °C) for 20 min during the gelation process. For cytotoxicity and ROS assays, 5 wt% GelMAL hydrogels were prepared as above, with primary hematopoietic stem and progenitor cells (HSPCs) resuspended at a density of 10^5^ cells/mL (cytotoxicity) or 5×10^5^ cells/mL (ROS) in the GelMAL precursor solution prior to dispensing into a Teflon mold (5 mm diameter, 20 µL) and addition of peptide crosslinker.

#### 2.3.3 GelMA hydrogels

A 5 wt% solution of GelMA was prepared in PBS. GelMA was functionalized following previously defined protocols [23, 69, 70] to match the degree of functionalization in GelMAL (∼80%). Lithium acylphosphinate photoinitiator (PI) was added to a final concentration of 0.1 wt% and 20 µL of the solution was dispensed into a Teflon mold (5 mm diameter, 20 µL) and exposed to 7.14 mW/cm^2^ UV light (λ = 365 nm) for 30 s. For cell response assays (cytotoxicity and ROS assay), HSPCs were resuspended at a density of 10^5^ cells/mL (cytotoxicity) or 5×10^5^ cells/mL (ROS) in the GelMA precursor solution prior to crosslinking.

### 2.4 Mechanical testing

Material properties of GelMAL with increasing gelatin content and crosslinkers of increasing molecular weight (DTT, peptide, PEG_1000_-dithiol) were investigated to determine the tunability of the system and appropriate matching of mechanics to the *in vivo* HSC niche. Hydrogels of GelMAL-DTT (3 and 5 wt%), GelMAL-peptide (5 wt%), and GelMAL-PEG (5 wt%) were swelled in PBS overnight. Compressive Young’s moduli of hydrogels (3 and 5 wt% GelMAL) were obtained from the stress–strain curve in unconfined compression using an Instron 5943 (Instron, Norwood, MA) as previously described [26, 71]. In brief, samples for mechanical testing were compressed at a rate of 0.1 mm/min and the moduli were calculated using a custom MATLAB code (Mathworks, Inc., Natick, MA). Raw compression data was smoothed with a moving average (n = 3) and contact with the hydrogel surface was established by 10 consecutive load outputs above 0.005 N. The moduli were calculated from a fit to the linear regime of 15% strain, with an initial offset of 2.5% strain. For swelling studies, 3 and 5 wt% GelMAL-DTT samples were hydrated in PBS for 24 h and weighed in air and in heptane using an Archimedes buoyancy test kit. Samples were then dehydrated in a vacuum oven at room temperature, and similarly weighed. The volume of samples was determined from the mass change in air and heptane, using the density of heptane to calculate volume displacement. The swelling ratio is defined as the ratio of the volume of the swollen hydrogel to the volume of the dehydrated hydrogel.

### 2.5 Cell isolation and cultures

All work involving primary cell extraction was conducted under approved animal welfare protocols (Institutional Animal Care and Use Committee, University of Illinois at Urbana-Champaign). Bone marrow cells were isolated from the crushed tibiae and femurs of C57BL/6 female mice, age 4-8 weeks (The Jackson Laboratory, Bar Harbor, ME). The bone marrow suspension was lysed in ACK lysis buffer for 2 min and then washed in PBS + 5% FBS (v/v) at 300 rcf for 10 min. Isolation of hematopoietic lineage negative was performed with EasySep™ Mouse Hematopoietic Progenitor Cell Isolation Kit (#19856, Stemcell Technologies, CA). For cytotoxicity studies, enriched lineage negative cells were further purified to obtain an enriched Lin-Sca-1+ c-kit+ (LSK) population using EasySep™ Mouse Sca1 and cKit Positive Selection kits (#18756 & #18757, Stemcell Technologies, CA). For ROS studies, a Lin^-^ Sca-1^+^ c-kit^+^ (LSK) hematopoietic stem and progenitor cell (HSPC) population was isolated using a BD FACS Aria II flow cytometer. LSK antibodies were supplied by eBioscience (San Diego, CA), and are as follows: APC-efluor780-conjugated c-kit (1:160, #47-1172-81), PE-conjugated Sca-1 (0.3:100, #12-5981-83), and Lin: FITC-conjugated CD5, B220, CD8a, CD11b (1:100, #11-0051-82, #11-0452-82, #11-0081-82, #11-0112-82), Gr-1 (1:400, #11-5931-82), and Ter-119 (1:200, #11-5921-82) [10, 65, 72].

Human bone marrow-derived mesenchymal stem cells (MSCs) (Lonza, Walkersville, MD) were cultured in DMEM supplemented with 10% MSC-qualified fetal bovine serum (FBS, Thermo Fisher Scientific, Waltham, MA), penicillin/streptomycin, and plasmocin prophylactic (Invivogen, San Diego, CA) to prevent mycoplasma contamination. The cells were used before passage 5 and were maintained in an incubator at 37 °C and 5% CO_2_.

### 2.6 Assessment of MSC Behavior in GelMAL Hydrogels

MSCs in 3 wt% and 5 wt% GelMAL were cultured for 72 h in DMEM supplemented with 10% MSC-qualified FBS, penicillin/streptomycin, and plasmocin prophylactic at 5% CO_2_ and 37 °C.

#### 2.6.1 Live/Dead Imaging

The Live/Dead Fixable Staining Kit (Promocell, Germany) was used to discriminate live and dead cells for imaging. Briefly, hydrogels were washed with PBS and incubated in 1 mL PBS containing 1 µL of Fixable Dead Cell Stain for 30 min with shaking. Subsequently, hydrogels were washed with PBS, fixed and permeabilized, and stained with Hoescht 33342 to visualize cell nuclei. Control hydrogels containing dead cells were prepared by fixing hydrogels in ice-cold 70% ethanol for 15 min (**Figure S1**). Hydrogels were imaged using DMi8 Yokogawa W1 spinning disk confocal microscope with a Hamamatsu EM-CCD digital camera (Leica Microsystems, Buffalo Grove, IL) by taking z-stacks at 10x magnification with a step size of 5 µm and a total depth of 200 µm. Three regions of interest were imaged per hydrogel. For analysis, background was subtracted from maximum intensity images using a rolling ball radius of five pixels in ImageJ (NIH, Bethesda, MD). Then, images were processed using CellProfiler to determine the percentage of live cells [73]. At least 1000 cells were imaged per hydrogel, with n = 3 hydrogels per group.

#### 2.6.2 F-Actin Staining and Imaging

Hydrogels were fixed with methanol-free formaldehyde and permeabilized using Tween 20. F-actin staining was performed using CytoPainter Phalloidin iFluor-633 Reagent (abcam, United Kindgom) for 1 h at RT. Hydrogels were washed and stained with Hoescht 33342 before imaging. Z-stacks were taken using a DMi8 Yokogawa W1 spinning disk confocal microscope with a Hamamatsu EM-CCD digital camera. Z-stacks had a step size of 2 µm and a total depth of 100 µm and were taken at 20x magnification.

### 2.7 Assessment of HSPC behavior in GelMAL hydrogels

#### 2.7.1 Cytotoxicity

5 wt% GelMAL-peptide and 5 wt% GelMA hydrogels containing 10^5^ LSK/mL were immediately placed in 500 µL SFEM media (#09650, Stemcell Technologies) supplemented with 100 ng/mL SCF (#250-03, Peprotech, Rocky Hill, NJ) and 1% PenStrep and cultured at 5% CO_2_ and 37 °C. At time points of 1 h and 48 h samples were tested for cytotoxicity via a commercially available LDH-Glo™ Cytotoxicity Assay (#J2380, Promega, Madison, WI). In brief, 10 µL conditioned media was diluted in LDH Storage Buffer (200 mM Tris-HCl, 10% glycerol, 1% BSA, pH 7.3) and reacted with LDH Detection Reagent on a shaker for 1 h at RT. Immediately following the reaction, the samples were moved to an opaque-walled 96-well plate in triplicate and luminescence was read using a BioTek Synergy HT Plate Reader and Gen5 Software (BioTek Instruments, Inc., Winooski, VT). Relative luminescence was calculated by subtracting the average values from a background control (media). Values of each sample are reported as the mean of each triplicate, and n = 4 (GelMAL-peptide) and n = 5 (GelMA).

#### 2.7.2 Reactive oxygen species

Purified LSK cells were encapsulated in either 5 wt% GelMAL (peptide) or GelMA (PI) at 5×10^5^ LSK/mL and cultured for 2 h at 5% CO_2_ and 37 °C in 500 µL SFEM media (#09650, Stemcell Technologies) supplemented with 100 ng/mL SCF (#250-03, Peprotech) and 1% PenStrep. After 2 h, hydrogels were resuspended in 5 μM CellROX Deep Red Reagent, for oxidative stress detection (#C10422, ThermoFisher Scientific) in media and incubated for 1 h. Hydrogels were washed in PBS for 5 min and fixed in 4% formaldehyde for 15 mins. Hydrogels were washed in PBS and stained with Hoescht 33342 (1:2000) before imaging. A negative control was created with no CellROX reagent (**Figure S2**). Z-stack images were captured using a Zeiss LSM 710 NLO (Zeiss, Oberkochen, Germany; 40x W objective). Maximum intensity images were processed using CellProfiler to quantify the mean CellROX fluorescence in each cell, with corrected fluorescence intensity reported as the fluorescence intensity with background (three reference spots) removed. A total of 4 mice were used to create n = 3 samples per hydrogel condition, with at least 10 cells per hydrogel (n = 30 cells per condition) and 3 reference spots per image.

### 2.9 Surface markers

Lin-murine bone marrow cells were resuspended in 100 µL degradation buffer (50 mM HEPES, 150 mM NaCl, 10 mM CaCl_2_, pH = 7.4) and exposed to either 100 Units Collagenase Type I, 100 Units of Collagenase Type IV (Worthington Biochemical, Lakewood, NJ), 10 µM SortaseA (#13101, ActiveMotif, Carlsbad, CA) with 15 mM GGG, and a control with buffer only. After 1 h at 37 °C, the degradation reaction was quenched with PBS + 5% FBS and centrifuged at 300 rcf x 10 min. The pellet was resuspended in PBS + 5% FBS for staining. After Fc blocking, cells were stained with surface marker antibodies and analyzed with Fluorescence-Assisted Cytometry (FACs) using a BD LSR Fortessa (BD Biosciences, San Jose, CA). Propidium Iodide (5:1000 µL) was used for dead cell exclusion. All samples were run for a minimum of 25,000 live events. Compensation was setup using AbC™ Total Antibody Compensation Bead Kit (#A10513, ThermoFisher). All antibodies were supplied by eBioscience (San Diego, CA) and are as follows: PE-conjugated CD150 (1:160, #12-1502-80), APC-conjugated CD48 (1:160, #17-0481-80), eFluor450-conjugated c-kit (4:500, #48-1171-80), PE-Cy7-conjugated Sca-1 (0.3:100, #25-5981-81), and Lin: FITC-conjugated CD5, B220, CD8a, CD11b (1:100, #11-0051-82, #11-0452-82, #11-0081-82, #11-0112-82), Gr-1 (1:400, #11-5931-82), and Ter-119 (1:200, #11-5921-82) [10, 65, 72]. A total of 4 mice were used to create n = 3 samples per degradation condition.

### 2.10 Statistical methods

Prior to significance testing, normality and equality of variance were tested with Shapiro-Wilks [74, 75] and Brown–Forsythe [76, 77] respectively at a significance level of 0.05. Surface markers were analyzed with a one-way ANOVA. Swelling ratio was analyzed with a Student’s T-test, and ROS studies were analyzed via Mann-Whitney U test. Elastic modulus studies were analyzed via a Welch-corrected ANOVA with Games-Howell post-hoc pairwise mean comparison. All statistical analysis was performed in R [78].

## 3. Results

### 3.1 Synthesis of GelMAL

Functionalized gelatin macromers were generated by decorating lysine amines along the gelatin macromer with maleimide functional groups *via* NHS ester-activated peptide coupling with maleimide reagent **M** (**Figure 1**), which was synthesized by reacting 3-maleimidopropionic acid with *N*-hydroxysuccinimide to give maleimide intermediate **M** in 75% yield (**Figure 1A**). Gelatin was mixed with 0.5 weight equivalent of **M** in a 1:1 (v/v) DMSO:water. The solvent (DMSO:water) minimized unwanted side reactions between lysine amines on gelatin and the maleimide group while maintaining sufficient reactivity between the amine and carboxyl. The resulting pH of 6.7 provided the appropriate balance between reactivity and stability of the maleimide, amine, and carboxyl groups; maleimide groups react selectively with thiols in a narrow reaction pH window (6.5-7.5), while higher pH (>8.5) favors primary amines reactions and increases the susceptibility of maleimides to hydrolysis [79]. The crude product was purified by dialysis and lyophilized to afford GelMAL as a light pink solid (**Figure 1B**).

**Figure 1.**
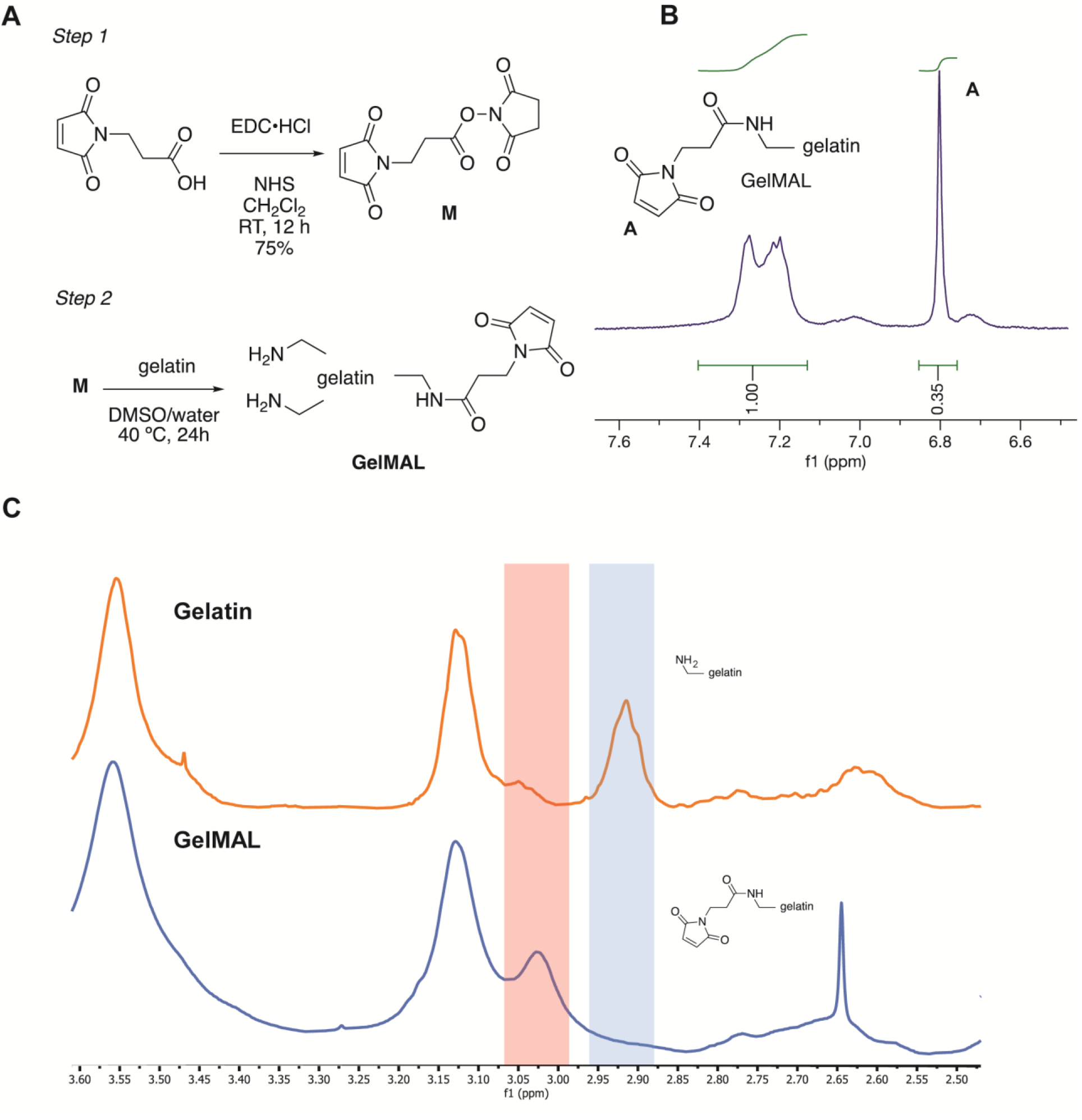
Chemical characterization of GelMAL. **A**. Two-step synthetic procedure (A) **B**. ^1^H NMR spectrum showing the chemical shift for the olefin protons on the maleimide ring in GelMAL **C**. Stacked ^1^H NMR spectra of unmodified gelatin and GelMAL showing shift in peak of hydrogen adjacent to nitrogen in lysine upon functionalization with maleimide.

### 3.2 Chemical characterization of GelMAL

The degree of functionalization (DOF) of GelMAL was defined as the percentage of amine groups from lysine and hydroxyl lysine that were functionalized with a maleimide group. The DOF was determined via ^1^H NMR spectroscopy as extensively reported previously by using the phenylalanine aromatic signal (7.5–6.9 ppm, integrating for 5 protons) as the reference integral and the decrease of the lysine *ε*-CH_2_ signal of nonmodified gelatin [54, 80] and GelMAL (2.95– 2.8 ppm). 2D NMR spectroscopy has been recently reported for gelatin and gelatin methacrylamide [81]; this method of analysis was performed on GelMAL to confirm the signals of the double bond protons of the maleimide group (6.8 ppm, integrating for 2 protons) (**Figure S3**). The measured ^1^H−^13^C-HSQC spectrum shows individual signals for each unique proton bonded to a ^13^C atom, providing more structural information than 1D ^1^H NMR spectroscopy [82], and serves as additional evidence that the lysine groups were functionalized (**Figure 1C)**. The integral of this signal with respect to the reference signal was also used to calculate the DOF of GelMAL and ranged from 70-85%.

### 3.3 Material properties of GelMAL hydrogel

Mechanical properties of GelMAL hydrogels were calculated as a function of increasing gelatin content and increasing molecular weight of crosslinker. Hydrogels of 3 or 5 wt% were crosslinked with either DTT (GelMAL-DTT), PEG_1000_-dithiol (GelMAL-PEG) or custom-made peptide (GelMAL-peptide). The peptide was designed to contain an LPRTG sequence to enable mammalian-inert cleavage by the SortaseA enzyme. The swelling ratio measures association of water and can be used to predict behavior of the hydrogel on multiple length scales (micro- and macro-properties) [83]. Volumetric swelling ratios (wet/dry volume) of 3 wt% and 5 wt% GelMAL-DTT were significantly different (p-value = 0.0381), with an increased ratio in the lower weight content hydrogel: 3 wt% = 26.5 ± 5.15 (n = 5), 5 wt% = 20.5 ± 2.85 (n = 6) (**Figure 2A**). Compression studies of GelMAL hydrogels (dia. =10 mm, DTT) at two different concentrations were performed to demonstrate the ability to control hydrogel stiffness. GelMAL-DTT hydrogels of 3 wt% vs. 5 wt% showed statistically significant (p = 0.018) differences in elastic modulus: E_3wt%_ = 1.81 ± 0.80 and E_5wt%_ = 4.58 ± 1.49 kPa (**Figure 2B**). Inserting a peptide or PEG_1000_ crosslinker into 5 wt% GelMAL led to a significant decrease in moduli compared to GelMAL-DTT, with E_PEG_ = 1.72 ± 0.60 (p-value = 0.018) and E_peptide_ = 1.49 ± 0.66 kPa (p-value = 0.010) respectively (**Figure 2B**). Increasing weight content led to an increase in elastic modulus and a decrease in swelling ratio, which is expected given the increased gelatin content.

**Figure 2.**
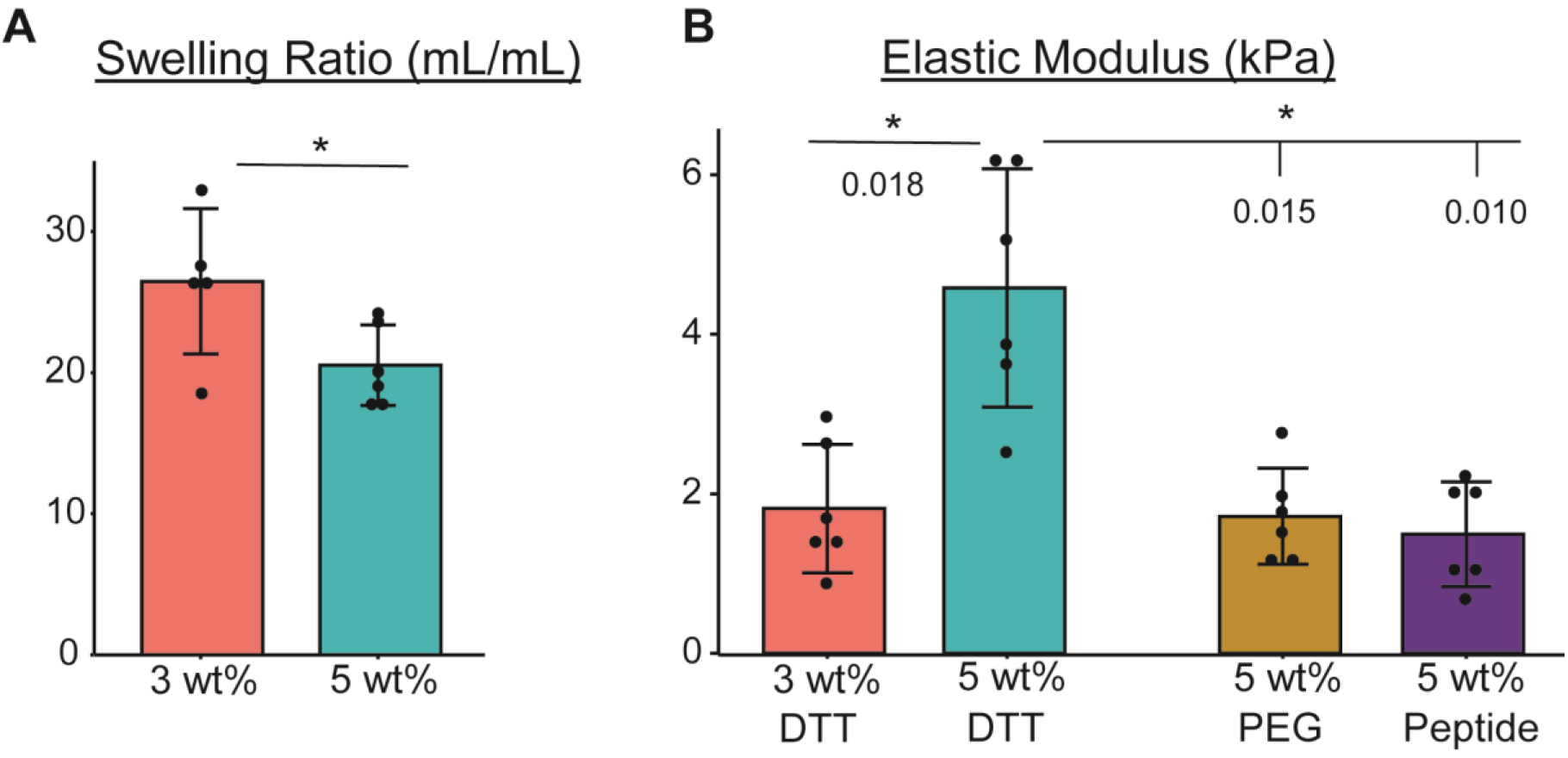
Biophysical characterization of GelMAL hydrogels. **A**. Volumetric swelling ratio of 3 and 5 wt% hydrogels. **n=5 and 6** respectively. (p-value = 0.0381). **B**. Elastic moduli were determined via unconfined mechanical compression of samples crosslinked via DTT (3 and 5 wt% GelMAL), PEG (5 wt% GelMAL), or peptide (5 wt% GelMAL). **n=6**. p-values displayed on graph. The bar height represents the mean, error bars are SD, and dots show the individual data points.

### 3.4 Cytotoxic response to GelMAL formation

HSC cytotoxicity in GelMAL hydrogels was compared to 5 wt% GelMA, which our lab has used extensively for HSC encapsulation [23, 26]. There was no detectable lactate dehydrogenase (LDH) in Lin-hematopoietic cell cultures 1 h post-encapsulation, setting a baseline (0) for the initial time point. While HSCs in GelMA hydrogels showed a slight increase in cytotoxicity after 48 h in culture, there was no significant difference (p-value = 0.275) in cytotoxicity between GelMA and GelMAL samples, with a relative luminescence of 5.58×10^4^ ± 5.65×10^4^ and 2.07×10^4^ ± 1.72×10^4^ (arbitrary units) respectively (**Figure S4**).

### 3.5 MSCs are viable in GelMAL hydrogels

To assess the ability for GelMAL hydrogels to support culture of multiple stem cell types, we encapsulated MSCs in hydrogels and performed live/dead viability (**Figure 3A**) and F-actin staining after 72 h. Qualitative and quantitative assessment of hydrogels stained using the Fixable Dead Cell Stain revealed that a majority of cells (> 95%) encapsulated in GelMAL hydrogels remained viable after long-term (72 h) culture (**Figure 3B**). Additionally, F-actin staining revealed that MSCs adopted spread morphologies in GelMAL hydrogels, suggesting the ability for the MSCs to stretch and bind to presented attachment sites (**Figure 3C**).

**Figure 3.**
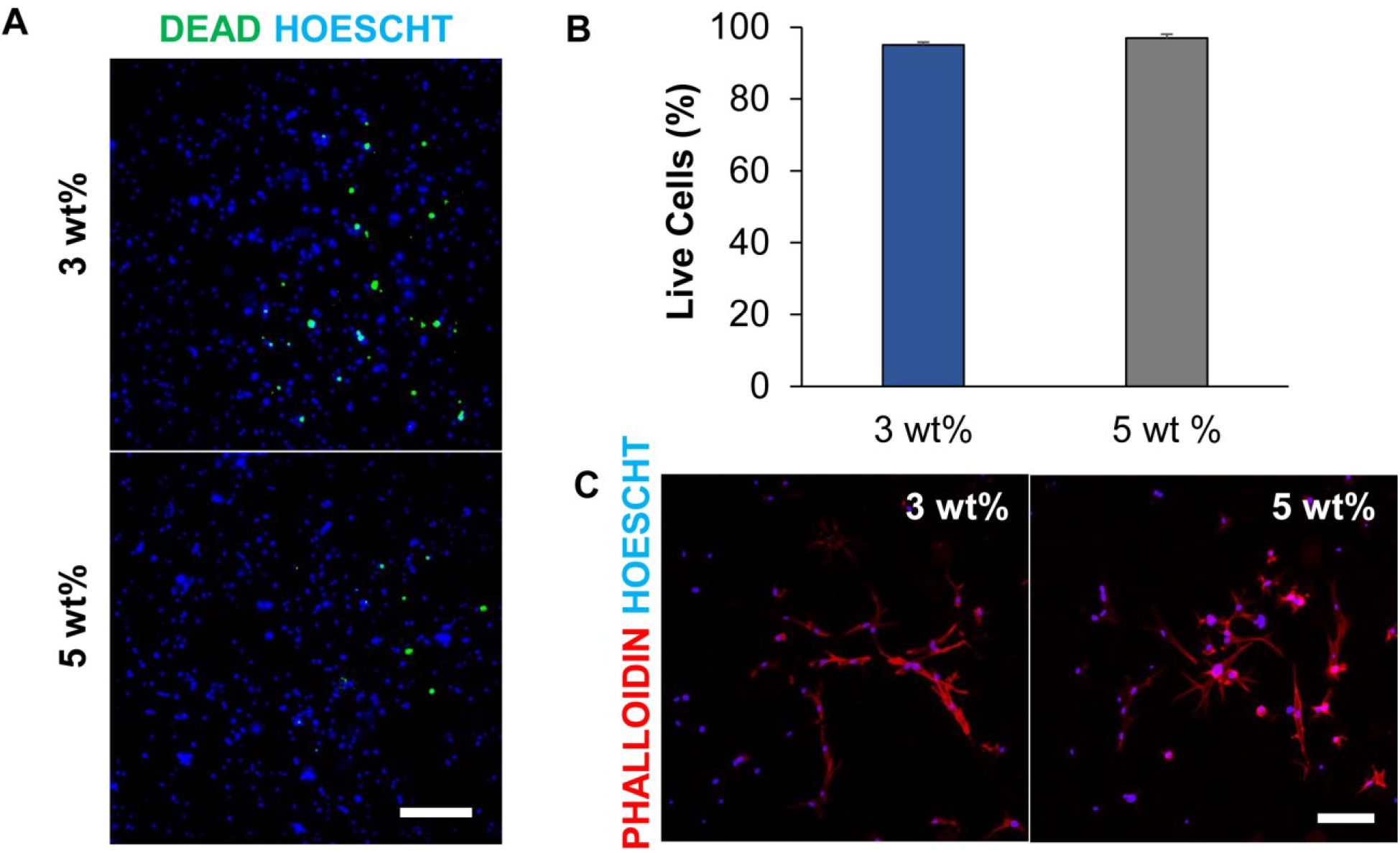
MSC behavior in GelMAL hydrogels. **A**. Representative images of hydrogels incubated with fixable live/dead reagent show minimal staining for dead cells. Scale bar = 200 µm **B**. Quantification of live cells reveals that the vast majority of encapsulated MSCs remain viable in 3 and 5 wt% GelMAL hydrogels. **C**. Phalloidin staining reveals that MSCs adopt a spread morphology in GelMAL hydrogels. Scale bar = 100 µm

### 3.6 Reactive oxygen species generation

Reactive oxygen species (ROS) generated during encapsulation of HSCs can artificially age and negatively influence stem cell fate decisions [64, 84, 85]. The use of photoinitiated crosslinking leads to increased concentration of radicals and ROS-mediated cytotoxicity propagate [56, 86, 87]. GelMAL, with insertable peptide crosslinks, is a route towards reducing ROS exposure of sensitive HSCs. Use of a dye that fluoresces when oxidized by reactive oxygen species showed increased oxidative stress on HSCs encapsulated in the presence of a photoinitiator (GelMA; **Figure 4A**). Comparison of total corrected fluorescence in cells encapsulated in conventional methacrylamide-functionalized gelatin (GelMA) versus maleimide crosslinked GelMAL-peptide (5 wt%) demonstrated a significant (p-value = 5.46 × 10^−5^) decrease in ROS uptake by HSCs in GelMAL (0.0290 ± 0.0137) compared to GelMA (0.0414 ± 0.00717; **Figure 4B**).

**Figure 4.**
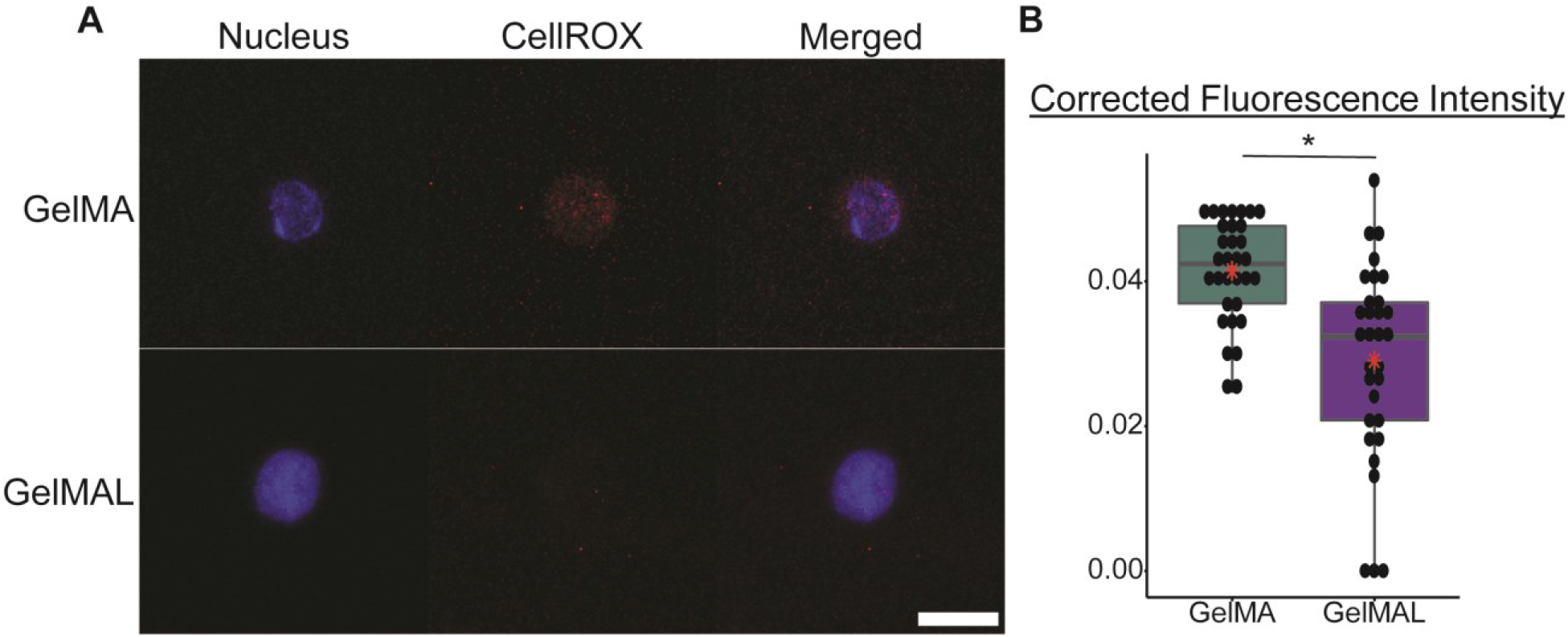
Fluorescent imaging of ROS. **A**. Representative images of an HSPC encapsulated in 5 wt% GelMA or GelMAL. Nucleus is stained with Hoescht. ROS are stained with CellROX (red). Images are auto-thresholded to Max/Min using ZEN lite (Zeiss Microscopy). Scale bar is 10 µm. **B**. Quantification of background corrected mean fluorescence intensity of CellROX dye in HSPCs encapsulated in 5 wt% GelMA (**n=32**) and GelMAL-peptide (**n=32**). Red crossbar represents mean, * indicates significant difference (p-value = 5.46 ×10^−5^).

### 3.7 Recovery of hematopoietic markers

We subsequently assessed the ability to analyze surface antigen expression of hematopoietic cells recovered after exposure to degradation enzymes used to degrade GelMAL-peptide (SortaseA) vs. GelMA (Collagenase I, Collagenase IV) hydrogels, with results compared to unexposed cells. SortaseA is an enzyme that specifically recognizes and cleaves the LPXTG motif, such as the LPRTG sequence found in the peptide crosslinker used in this study. Common surface markers of hematopoietic stem cells were analyzed after exposure to degradation enzymes (**Table 1**). Of the 5 surface markers tested, 3 were sensitive to the degradation scheme used. CD150 and c-Kit expression were both significantly lower in Collagenase treated samples compared to SortaseA, with p-value < 0.0001 for each condition. **Figure 5** demonstrates a representative LSK isolation scheme where Live events are isolated from the Singlet events, followed by gating on the lineage negative population (**Figure 5A**) and subsequently the Sca1+ and c-Kit+ subpopulation (**Figure 5B**). Gating on lineage-committed negative cells (Lin-) is the first step to purifying hematopoietic stem cells (**Figure 5A**). There was a significant increase in Lin-cells in samples exposed to Collagenase Type I (78.4 ± 0.495 %) or Collagenase Type IV (77.0 ± 0.739 %) compared to SortaseA (54.5 ± 11.4 %) and control (60.0 ± 1.41 %) cells. While the overall percentage of Lin-cells increased with collagenase exposure, there was a decrease in the LSK (Lin-Sca-1+ c-Kit+) hematopoietic stem and progenitor cell population obtained after collagenase exposure. We observed no differences in the fraction of identified LSK cells between control and SortaseA (5.06 ± 0.207% of all cells) exposed cells. However, exposure to Collagenase Type I (0.0398 ± 0.0041 %) or Collagenase Type IV (0.256 ± 0.0464 %) led to a significant decrease in overall identified LSK cells (p-values < 3.8×10^−9^ compared to SortaseA exposure).

**Table 1.**
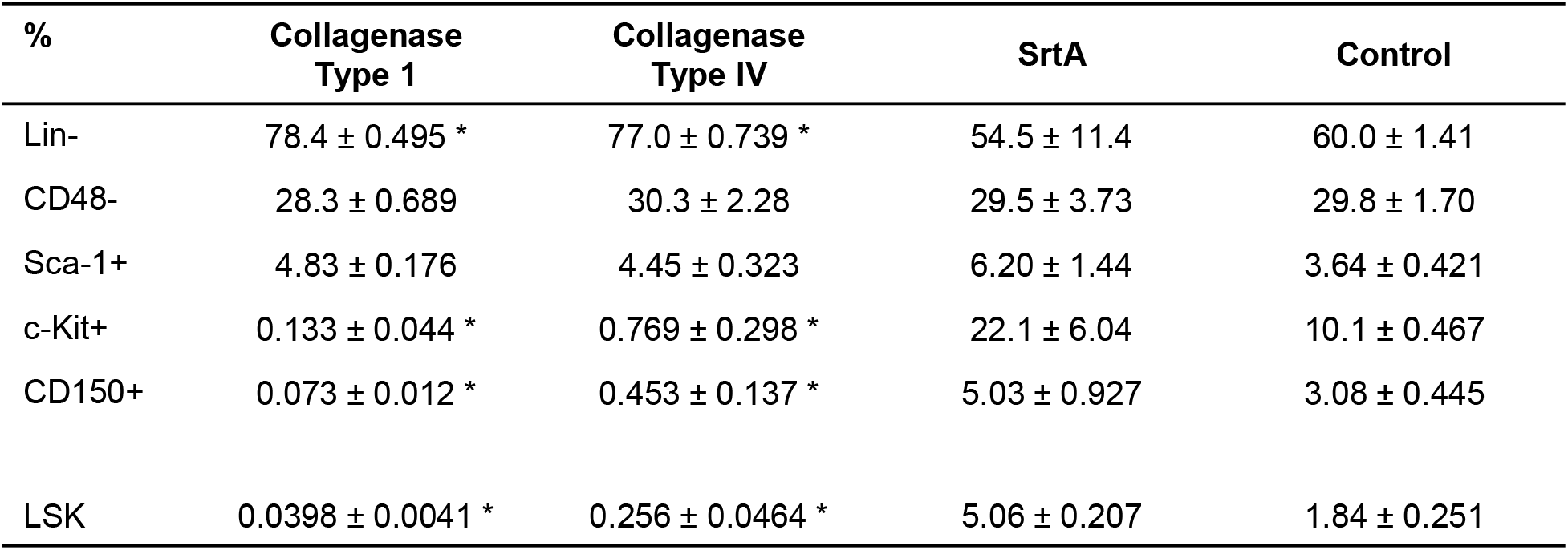
Percent of live cells after exposure to degradative enzyme. * Indicates significance compared to SrtA with p-value<0.05

| % | Collagenase<br>Type 1 | Collagenase<br>Type IV | SrtA | Control |
| --- | --- | --- | --- | --- |
| Lin- | 78.4 ± 0.495 * | 77.0 ± 0.739 * | 54.5 ± 11.4 | 60.0 ± 1.41 |
| CD48- | 28.3 ± 0.689 | 30.3 ± 2.28 | 29.5 ± 3.73 | 29.8 ± 1.70 |
| Sca-1+ | 4.83 ± 0.176 | 4.45 ± 0.323 | 6.20 ± 1.44 | 3.64 ± 0.421 |
| c-Kit+ | 0.133 ± 0.044 * | 0.769 ± 0.298 * | 22.1 ± 6.04 | 10.1 ± 0.467 |
| CD150+ | 0.073 ± 0.012 * | 0.453 ± 0.137 * | 5.03 ± 0.927 | 3.08 ± 0.445 |
| LSK | 0.0398 ± 0.0041 * | 0.256 ± 0.0464 * | 5.06 ± 0.207 | 1.84 ± 0.251 |

**Figure 5.**
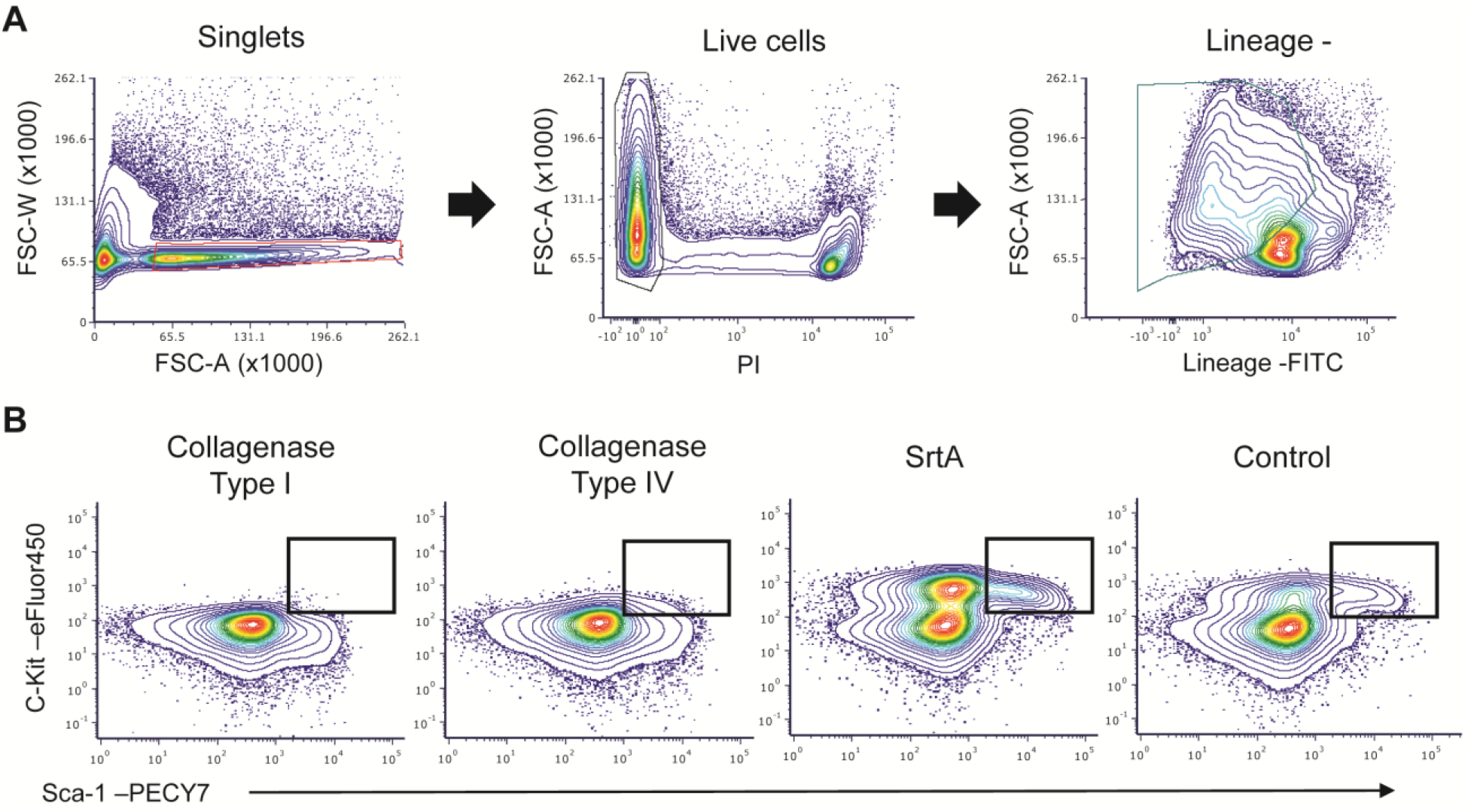
Representative gating strategy. **A**. Representative FACs graphs showing gating hierarchy of Singlet events to dead cell exclusion to gating on lineage negative events. **B**. Identification of LSK (Lin-Sca-1+ c-Kit+) cells after exposure to degradative enzyme. All graphs show Lineage negative events identified using the hierarchy listed in A.

## 4. Discussion

Material systems for stem cell culture are an important step in developing methods for expansion or interrogation of stem cell biology. Commonly used biomaterials, such as PEG, require additional insertion of cell-interaction sites or expose encapsulated stem cells to damaging reactive oxygen species (ROS) during crosslinking. Gelatin is a commonly used material for stem cell culture, and it has previously been applied to stem cell systems due to its poroelastic (modulus, diffusion) tunability and its inherent cell interaction sites (e.g. the cell-adhesion site RGD). Although comprised mainly of glycine (∼33%) [88], the free amines present in lysine groups along the backbone of gelatin offers an avenue to create a library of chemical modifications. We report a two-step synthesis process that leverages easily accessible and standard N-hydroxysuccinimide (NHS) chemistries to functionalize free amines on the gelatin macromer with pendulant maleimides with a high degree of functionalization (70 - 80%). This effort, while inspired by recent development of maleimide-functionalized PEG hydrogels [89-91] is to our knowledge the first report of the modification of lysine groups within a gelatin macromer with maleimide groups.

The tunability of the GelMAL hydrogels system provides biophysical features that recapitulate the stem cell microenvironment. Using HSCs as a model system, the mechanical properties (elastic modulus, swelling ratio) of GelMAL hydrogels were tuned to match the *in vivo* HSC microenvironment (0.25 – 24 kPa) [92] which has previously been shown to maintain an HSC population *in vitro* [23]. Increasing gelatin content (3 to 5 wt%) led to 2.5-fold increase in modulus in DTT crosslinked GelMAL and a decrease in swelling ratio. The flexibly of the system offers a choice of crosslinkers, allowing for a range of modulus without loss of gelatin content, which possesses the required cell-interaction motifs essential for stem cell culture. By altering the choice of crosslinker from a small molecule (DTT, 154.25 g/mol) to a larger molecule (PEG, 1000 g/mol; peptide, 1405.58 g/mol) there was an associated 2.5-fold decrease in modulus of 5 wt% hydrogels.

The dithiol-based crosslinking of GelMAL offers versatility to orthogonally modulate a range of biophysical properties without adverse cytotoxic response. Notably, we described the potential for insertion of a cleavable peptide (GCRD-LPRTG-DRCG), capped by two thiol-presenting cysteines. The inserted peptide can be cleaved using the bacterial-based enzymatic reaction of Sortase A (SrtA), which is highly specific to the LPXTG motif [68, 93] and is a standard in protein engineering [94-96]. Gelatin-based hydrogels are commonly degraded with mammalian collagenases that cleave along the gelatin backbone. However, collagenase degradation is promiscuous and can cause degradation of essential biomarkers for hemopoietic activity. Collagenase Type I and the less tryptic Collagenase Type IV [97] can cleave or cause internalization of surface markers required for stem cell characterization. The insertion of a SrtA-cleavable peptide during crosslinking enables degradation without injury to surface markers enabling accurate quantification of hematopoietic populations and biomolecular activity. Surface markers present on hematopoietic cells are essential for immunophenotyping assays that are the backbone of hematopoietic identification and isolation. We show that while exposure to SrtA preserved commonly used markers of hematopoietic stem cells (LSK), commonly used collagenase enzymes led to a loss in surface marker fidelity. This may be particularly valuable for rapid isolation and analysis of rare cell populations as well as to adapt approaches used to examine the complex secretome generated by encapsulated cells within synthetic PEG hydrogels now within gelatin hydrogels.

Importantly, dithiol-based crosslinking of a maleimide-functionalized gelatin allows for successful encapsulation of primary murine hematopoietic stem cells without the introduction of confounding reactive oxygen species (ROS). By avoiding free radical-initiated crosslinking schemes, the production of ROS during encapsulation can be mitigated, as evidenced by the significantly decreased ROS uptake in HSCs encapsulated in GelMAL. Exposure of HSCs to ROS leads to a decrease in lifespan and alteration of self-renewal and differentiation pathways [62-64]. Additionally, the dithiolated peptide crosslinker showed no cytotoxic effect on sensitive HSCs even after extended culture. Similarly, MSCs encapsulated in GelMAL (DTT-crosslinked) have high long-term viability and the spreading morphology of MSCs within GelMAL confirms local cell-matrix interactions and paves the way for future studies examining dynamic reciprocity: the synergistic interactions between matrix and different cell types [98, 99]. Taken together, these data suggest that material properties of GelMAL can be tuned to mimic the *in vivo* microenvironment in a manner that reduces concerns associated with crosslinking-mediate ROS production and using degradation schemes less likely to cleave cell surface antigens or secreted biomolecules.

The reported GelMAL hydrogel provides important advantages for the development of *ex vivo* culture platforms for expansion and analysis of (stem) cells in multidimensional hydrogels. The GelMAL hydrogel provides a route to tune biophysical properties, incorporate intrinsic cell-interaction motifs, and add orthogonal cleavable crosslinks. The GelMAL hydrogel successfully encapsulated two disparate adult stem cell populations commonly localized within the bone marrow (HSCs and MSCs) without ROS production and retains high levels of cell viability. We also show the dithiol crosslinking approach enables insertion of a peptide that can be cleaved with a mammalian-inert proteolytic enzyme. This is an important step in furthering investigations into stem cell fate decisions with unhindered recovery of vital surface markers. GelMAL hydrogels may be particularly valuable in future studies of reciprocal signaling between multiple cell populations. The potential for rapid, target degradation of GelMAL via a Sortase-mediated reaction is critical for such studies where radical propagation or cell isolation process would reduce cell viability and hinder quantitative assessment of cell phenotype.

## Acknowledgments

The authors would like to acknowledge Dr. Barbara Pilas and Barbara Balhan of the Roy J. Carver Biotechnology Center (Flow Cytometry Facility, UIUC) for assistance with bone marrow cell isolation and flow cytometry. Research reported in this publication was supported by the National Institute of Diabetes and Digestive and Kidney Diseases of the National Institutes of Health under Award Numbers R01 DK099528 (B.A.C.H) and F31 DK117514 (A.E.G.), the National Cancer Institute of the National Institutes of Health under Award Number R01 CA197488 (B.A.C.H.), as well as by the National Institute of Biomedical Imaging and Bioengineering of the National Institutes of Health under Award Numbers R21 EB018481 (B.A.C.H.) and T32 EB019944 (A.E.G.). The authors also acknowledge funding from the National Science Foundation Graduate Research Fellowship (DGE 1144245, M.T.N.). The content is solely the responsibility of the authors and does not necessarily represent the official views of the NIH or the NSF. The authors are also grateful for additional funding provided by the Department of Chemical & Biomolecular Engineering and the Institute for Genomic Biology at the University of Illinois at Urbana-Champaign.

## Supplemental Figures

**Figure S1.**
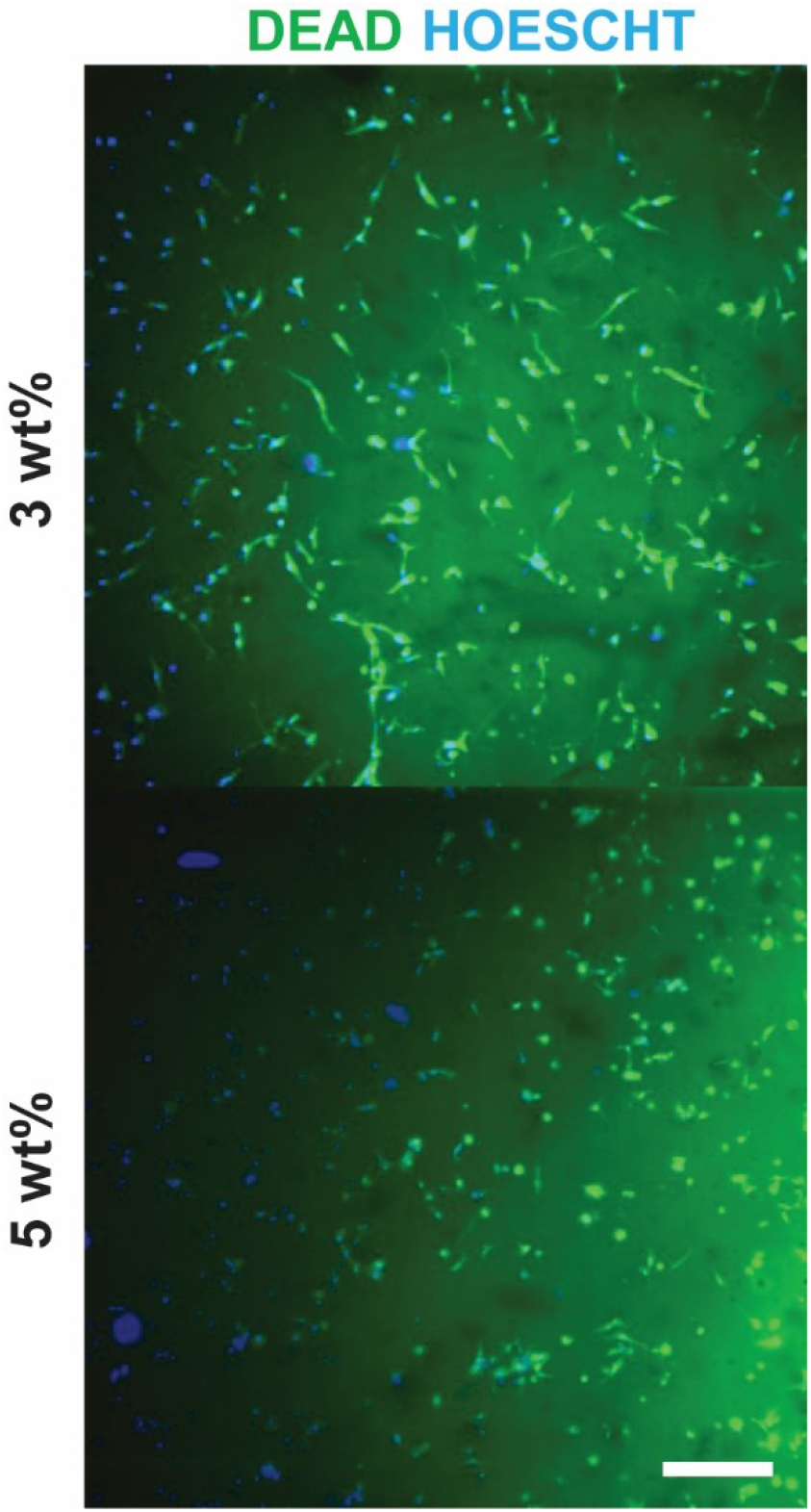
Hydrogels containing MSCs were fixed with ice-cold ethanol to generate dead cell controls for fixable live/dead staining. Scale bar = 200 µm

**Figure S2.**
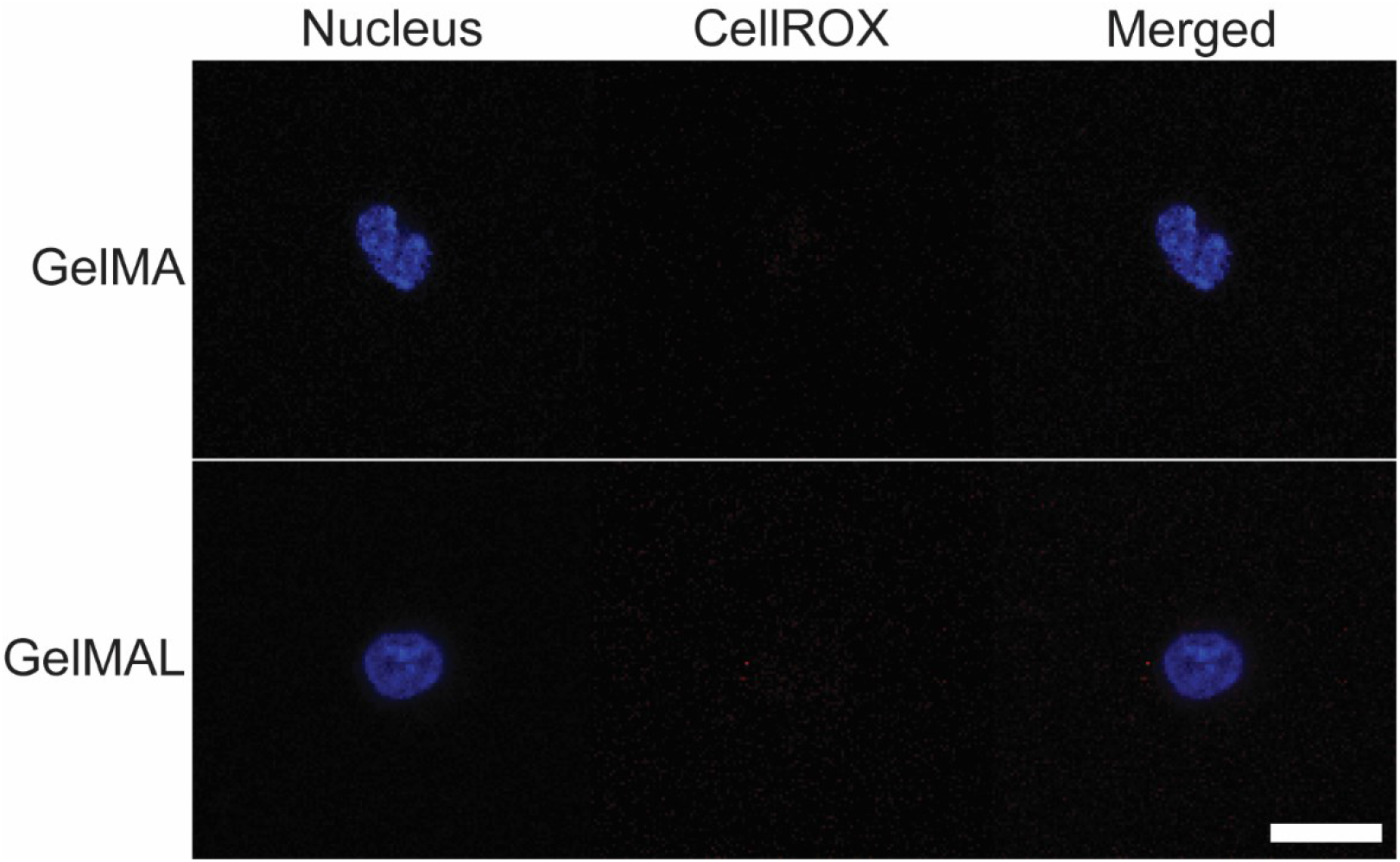
Negative controls of GelMA and GelMAL with no CellROX staining. Scale bar = 10 µm

**Figure S3.**
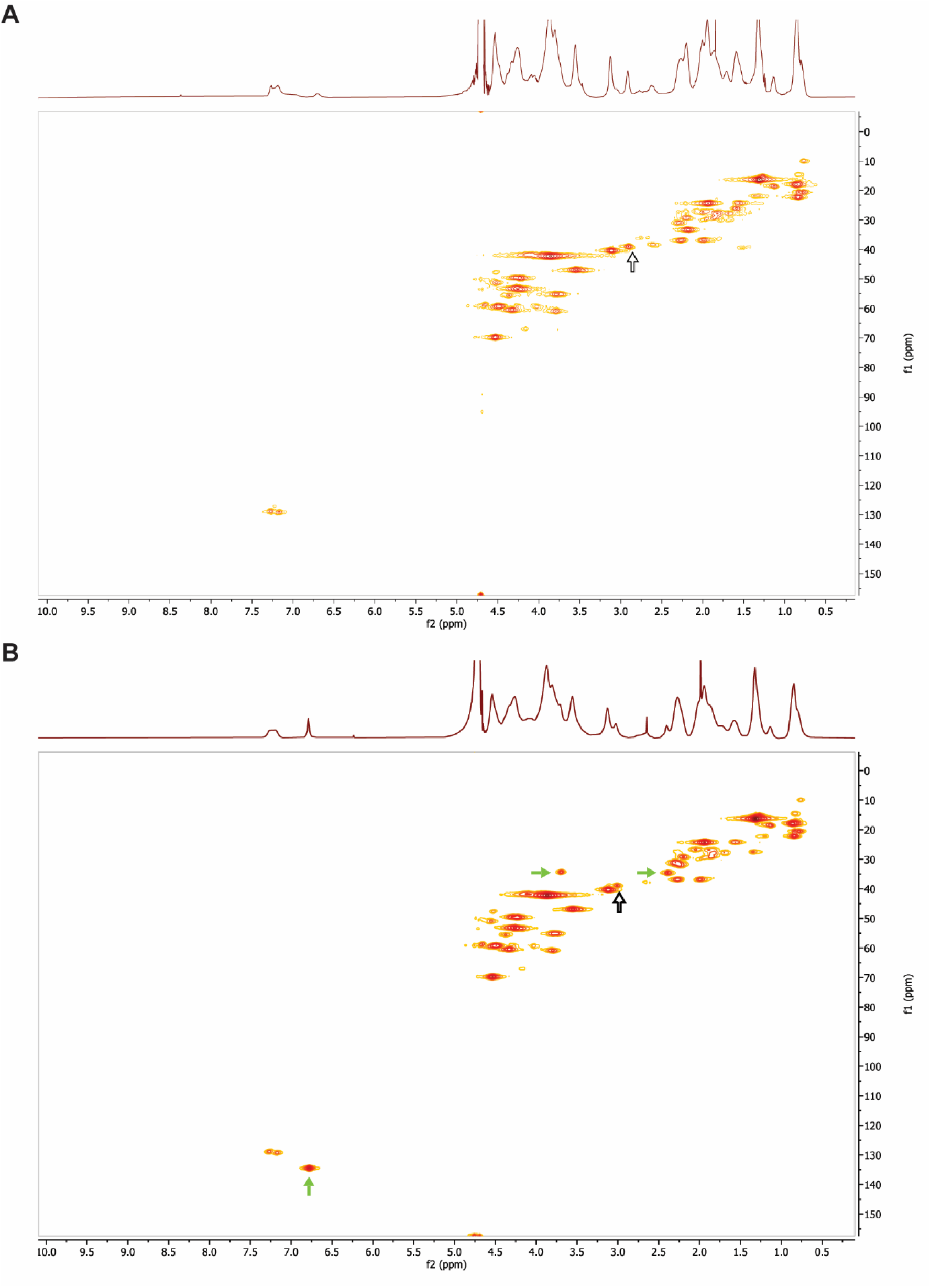
^1^H−^13^C-HSQC 2D NMR Spectroscopy. A. Spectrum of unmodified gelatin. B. Spectrum of maleimide-functionalized gelatin. Filled arrows point to emergence of peaks from maleimide. Empty arrows point to a shift from 2.89 to 3.02 ppm of the modified lysine. X-axis is ^1^H and y-axis is ^13^C.

**Figure S4.**
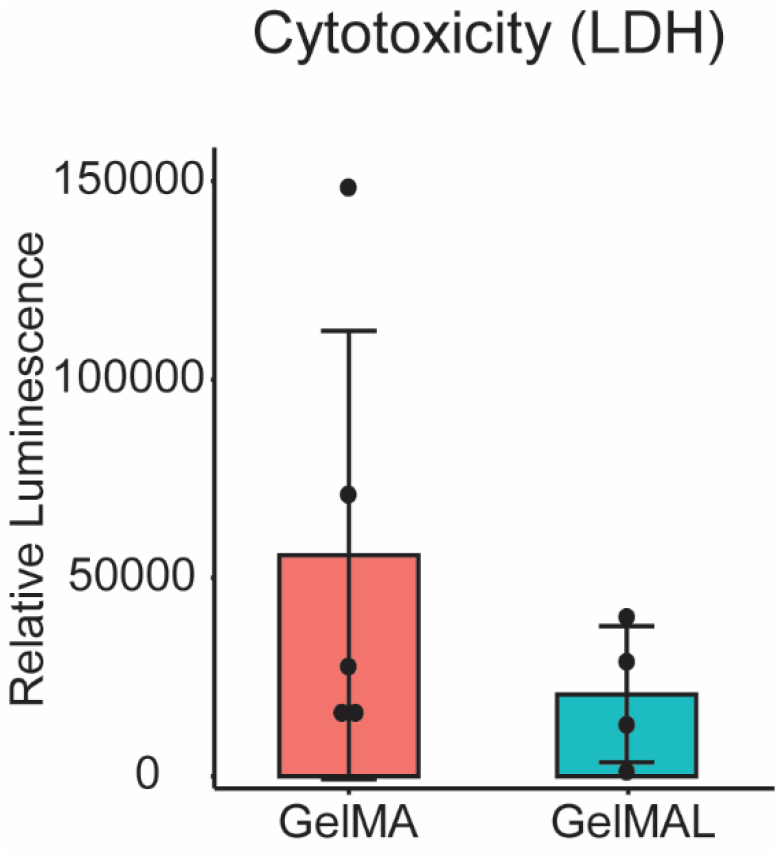
Relative luminescence of LDH present in GelMA and GelMAL after 48hr culture. Relative luminescence with background subtracted using a media control.

## References

[1] S.J. Morrison, N.M. Shah, D.J. Anderson, Regulatory mechanisms in stem cell biology, Cell 88(3) (1997) 287–98.

[2] R.R. Nadig, Stem cell therapy - Hype or hope? A review, J Conserv Dent 12(4) (2009) 131–8.

[3] A. Usas, J. Huard, Muscle-derived stem cells for tissue engineering and regenerative therapy, Biomaterials 28(36) (2007) 5401–6.

[4] K.H. Schuleri, A.J. Boyle, J.M. Hare, Mesenchymal stem cells for cardiac regenerative therapy, Handb Exp Pharmacol (180) (2007) 195–218.

[5] A. Mendelson, P.S. Frenette, Hematopoietic stem cell niche maintenance during homeostasis and regeneration, Nat Med 20(8) (2014) 833–46.

[6] M.F. Pittenger, D.E. Discher, B.M. Peault, D.G. Phinney, J.M. Hare, A.I. Caplan, Mesenchymal stem cell perspective: cell biology to clinical progress, NPJ Regen Med 4 (2019) 22.

[7] C. Brown, C. McKee, S. Bakshi, K. Walker, E. Hakman, S. Halassy, D. Svinarich, R. Dodds, C.K. Govind, G.R. Chaudhry, Mesenchymal stem cells: Cell therapy and regeneration potential, J Tissue Eng Regen Med 13(9) (2019) 1738–1755.

[8] N.S. Hwang, C. Zhang, Y.S. Hwang, S. Varghese, Mesenchymal stem cell differentiation and roles in regenerative medicine, Wiley Interdiscip Rev Syst Biol Med 1(1) (2009) 97–106.

[9] S. Doulatov, F. Notta, E. Laurenti, J.E. Dick, Hematopoiesis: a human perspective, Cell Stem Cell 10(2) (2012) 120–36.

[10] L. Yang, D. Bryder, J. Adolfsson, J. Nygren, R. Mansson, M. Sigvardsson, S.E. Jacobsen, Identification of Lin(-)Sca1(+)kit(+)CD34(+)Flt3-short-term hematopoietic stem cells capable of rapidly reconstituting and rescuing myeloablated transplant recipients, Blood 105(7) (2005) 2717–23.

[11] A. D’Souza, C. Fretham, S.J. Lee, M. Arora, J. Brunner, S. Chhabra, S. Devine, M. Eapen, M. Hamadani, P. Hari, M.C. Pasquini, W. Perez, R.A. Phelan, M.L. Riches, J.D. Rizzo, W. Saber, B.E. Shaw, S.R. Spellman, P. Steinert, D.J. Weisdorf, M.M. Horowitz, Current Use of and Trends in Hematopoietic Cell Transplantation in the United States, Biol Blood Marrow Transplant 26(8) (2020) e177–e182.

[12] E.A. Copelan, Hematopoietic stem-cell transplantation, N Engl J Med 354(17) (2006) 1813–26.

[13] S. Nikiforow, J. Ritz, Dramatic Expansion of HSCs: New Possibilities for HSC Transplants?, Cell Stem Cell 18(1) (2016) 10–2.

[14] V. Rocha, E. Gluckman, r. Eurocord-Netcord, B. European, g. Marrow Transplant, Improving outcomes of cord blood transplantation: HLA matching, cell dose and other graft-and transplantation-related factors, Br J Haematol 147(2) (2009) 262–74.

[15] A.T.L. Lam, S. Reuveny, S.K. Oh, Human mesenchymal stem cell therapy for cartilage repair: Review on isolation, expansion, and constructs, Stem Cell Res 44 (2020) 101738.

[16] R.A. Marklein, J.A. Burdick, Controlling stem cell fate with material design, Adv Mater 22(2) (2010) 175–89.

[17] C. Cha, W.B. Liechty, A. Khademhosseini, N.A. Peppas, Designing biomaterials to direct stem cell fate, ACS Nano 6(11) (2012) 9353–8.

[18] Z. Liu, M. Tang, J. Zhao, R. Chai, J. Kang, Looking into the Future: Toward Advanced 3D Biomaterials for Stem-Cell-Based Regenerative Medicine, Adv Mater 30(17) (2018) e1705388.

[19] R.H. Li, D.H. Altreuter, F.T. Gentile, Transport characterization of hydrogel matrices for cell encapsulation, Biotechnol Bioeng 50(4) (1996) 365–73.

[20] R.J. McMurtrey, Roles of Diffusion Dynamics in Stem Cell Signaling and Three-Dimensional Tissue Development, Stem Cells Dev 26(18) (2017) 1293–1303.

[21] S.Y. Shvartsman, H.S. Wiley, W.M. Deen, D.A. Lauffenburger, Spatial Range of Autocrine Signaling: Modeling and Computational Analysis, Biophysical Journal 81(4) (2001) 1854–1867.

[22] J. Hansing, J.R. Duke, 3rd, E.B. Fryman, J.E. DeRouchey, R.R. Netz, Particle Diffusion in Polymeric Hydrogels with Mixed Attractive and Repulsive Interactions, Nano Lett 18(8) (2018) 5248–5256.

[23] A.E. Gilchrist, S. Lee, Y. Hu, B.A.C. Harley, Soluble Signals and Remodeling in a Synthetic Gelatin-Based Hematopoietic Stem Cell Niche, Adv Healthc Mater 8(20) (2019) e1900751.

[24] M.T. Ngo, V.R. Barnhouse, A.E. Gilchrist, C.J. Hunter, J.N. Hensold, B.A.C. Harley, Hydrogels containing gradients in vascular density reveal dose-dependent role of angiocrine cues on stem cell behavior, (under review, 2021).

[25] B.P. Mahadik, N.A. Bharadwaj, R.H. Ewoldt, B.A. Harley, Regulating dynamic signaling between hematopoietic stem cells and niche cells via a hydrogel matrix, Biomaterials 125 (2017) 54–64.

[26] A.E. Gilchrist, B.A.C. Harley, Connecting secretome to hematopoietic stem cell phenotype shifts in an engineered bone marrow niche, Integr Biol (Camb) 12(7) (2020).

[27] E. Muller, W. Wang, W. Qiao, M. Bornhauser, P.W. Zandstra, C. Werner, T. Pompe, Distinguishing autocrine and paracrine signals in hematopoietic stem cell culture using a biofunctional microcavity platform, Sci Rep 6 (2016) 31951.

[28] M.P. Lutolf, H.M. Blau, Artificial stem cell niches, Adv Mater 21(32-33) (2009) 3255–68.

[29] T. Li, Y. Wu, TParacrine molecules of mesenchymal stem cells for hematopoietic stem cell niche, Bone Marrow Res 2011 (2011) 353878.

[30] B.P. Mahadik, S. Pedron Haba, L.J. Skertich, B.A. Harley, The use of covalently immobilized stem cell factor to selectively affect hematopoietic stem cell activity within a gelatin hydrogel, Biomaterials 67 (2015) 297–307.

[31] J.S. Choi, B.A. Harley, Marrow-inspired matrix cues rapidly affect early fate decisions of hematopoietic stem and progenitor cells, Sci Adv 3(1) (2017) e1600455.

[32] J. Jiang, E.T. Papoutsakis, Stem-cell niche based comparative analysis of chemical and nano- mechanical material properties impacting ex vivo expansion and differentiation of hematopoietic and mesenchymal stem cells, Adv Healthc Mater 2(1) (2013) 25–42.

[33] J.A. LaIuppa, T.A. McAdams, E.T. Papoutsakis, W.M. Miller, Culture materials affect ex vivo expansion of hematopoietic progenitor cells, J Biomed Mater Res 36(3) (1997) 347–59.

[34] M.L. Cuchiara, S. Coskun, O.A. Banda, K.L. Horter, K.K. Hirschi, J.L. West, Bioactive poly(ethylene glycol) hydrogels to recapitulate the HSC niche and facilitate HSC expansion in culture, Biotechnol Bioeng 113(4) (2016) 870–81.

[35] A. Raic, L. Rodling, H. Kalbacher, C. Lee-Thedieck, Biomimetic macroporous PEG hydrogels as 3D scaffolds for the multiplication of human hematopoietic stem and progenitor cells, Biomaterials 35(3) (2014) 929–40.

[36] D.S.W. Benoit, A.R. Durney, K.S. Anseth, The effect of heparin-functionalized PEG hydrogels on three-dimensional human mesenchymal stem cell osteogenic differentiation, Biomaterials 28(1) (2007) 66–77.

[37] S.Q. Liu, R. Tay, M. Khan, P.L. Rachel Ee, J.L. Hedrick, Y.Y. Yang, Synthetic hydrogels for controlled stem cell differentiation, Soft Matter 6(1) (2010) 67–81.

[38] K. Yue, G. Trujillo-de Santiago, M.M. Alvarez, A. Tamayol, N. Annabi, A. Khademhosseini, Synthesis, properties, and biomedical applications of gelatin methacryloyl (GelMA) hydrogels, Biomaterials 73 (2015) 254–71.

[39] M. Nikkhah, M. Akbari, A. Paul, A. Memic, A. Dolatshahi-Pirouz, A. Khademhosseini, Gelatin-Based Biomaterials For Tissue Engineering And Stem Cell Bioengineering, Biomaterials from Nature for Advanced Devices and Therapies2016, pp. 37–62.

[40] A.B. Bello, D. Kim, D. Kim, H. Park, S.H. Lee, Engineering and Functionalization of Gelatin Biomaterials: From Cell Culture to Medical Applications, Tissue Eng Part B Rev 26(2) (2020) 164–180.

[41] J.W. Nichol, S.T. Koshy, H. Bae, C.M. Hwang, S. Yamanlar, A. Khademhosseini, Cell-laden microengineered gelatin methacrylate hydrogels, Biomaterials 31(21) (2010) 5536–44.

[42] B.J. Klotz, D. Gawlitta, A. Rosenberg, J. Malda, F.P.W. Melchels, Gelatin-Methacryloyl Hydrogels: Towards Biofabrication-Based Tissue Repair, Trends Biotechnol 34(5) (2016) 394–407.

[43] X.S. Jiang, C. Chai, Y. Zhang, R.X. Zhuo, H.Q. Mao, K.W. Leong, Surface-immobilization of adhesion peptides on substrate for ex vivo expansion of cryopreserved umbilical cord blood CD34+ cells, Biomaterials 27(13) (2006) 2723–32.

[44] S.M. Dellatore, A.S. Garcia, W.M. Miller, Mimicking stem cell niches to increase stem cell expansion, Curr Opin Biotechnol 19(5) (2008) 534–40.

[45] C.N. Salinas, K.S. Anseth, The influence of the RGD peptide motif and its contextual presentation in PEG gels on human mesenchymal stem cell viability, J Tissue Eng Regen Med 2(5) (2008) 296–304.

[46] J. Poppe, Gelatin, Thickening and Gelling Agents for Food1997, pp. 144–168.

[47] M. Djabourov, J. Leblond, P. Papon, Gelation of aqueous gelatin solutions. I. Structural investigation, Journal de Physique 49(2) (1988) 319–332.

[48] B. Brodsky, J.A. Werkmeister, J.A.M. Ramshaw, Collagens and Gelatins, Biopolymers Online2005.

[49] C.M. Ofner, 3rd, W.A. Bubnis, Chemical and swelling evaluations of amino group crosslinking in gelatin and modified gelatin matrices, Pharm Res 13(12) (1996) 1821–7.

[50] E. Schacht, B. Bogdanov, A.V.D. Bulcke, N. De Rooze, Hydrogels prepared by crosslinking of gelatin with dextran dialdehyde, Reactive and Functional Polymers 33(2-3) (1997) 109–116.

[51] J.A. Benton, C.A. DeForest, V. Vivekanandan, K.S. Anseth, Photocrosslinking of gelatin macromers to synthesize porous hydrogels that promote valvular interstitial cell function, Tissue Eng Part A 15(11) (2009) 3221–30.

[52] S. Pedron, B.A. Harley, Impact of the biophysical features of a 3D gelatin microenvironment on glioblastoma malignancy, J Biomed Mater Res A 101(12) (2013) 3404–15.

[53] E. Kaemmerer, F.P. Melchels, B.M. Holzapfel, T. Meckel, D.W. Hutmacher, D. Loessner, Gelatine methacrylamide-based hydrogels: an alternative three-dimensional cancer cell culture system, Acta Biomater 10(6) (2014) 2551–62.

[54] H. Shirahama, B.H. Lee, L.P. Tan, N.J. Cho, Precise Tuning of Facile One-Pot Gelatin Methacryloyl (GelMA) Synthesis, Sci Rep 6 (2016) 31036.

[55] X. Zhao, Q. Lang, L. Yildirimer, Z.Y. Lin, W. Cui, N. Annabi, K.W. Ng, M.R. Dokmeci, A.M. Ghaemmaghami, A. Khademhosseini, Photocrosslinkable Gelatin Hydrogel for Epidermal Tissue Engineering, Adv Healthc Mater 5(1) (2016) 108–18.

[56] A.K. Nguyen, P.L. Goering, R.K. Elespuru, S. Sarkar Das, R.J. Narayan, The Photoinitiator Lithium Phenyl (2,4,6-Trimethylbenzoyl) Phosphinate with Exposure to 405 nm Light Is Cytotoxic to Mammalian Cells but Not Mutagenic in Bacterial Reverse Mutation Assays, Polymers (Basel) 12(7) (2020).

[57] D.Q. Tan, T. Suda, Reactive Oxygen Species and Mitochondrial Homeostasis as Regulators of Stem Cell Fate and Function, Antioxid Redox Signal 29(2) (2018) 149–168.

[58] E. Peerani, J.B. Candido, D. Loessner, Cell Recovery of Hydrogel-Encapsulated Cells for Molecular Analysis, Methods Mol Biol 2054 (2019) 3–21.

[59] M. Abuzakouk, C. Feighery, C. O’Farrelly, Collagenase and Dispase enzymes disrupt lymphocyte surface molecules, Journal of Immunological Methods 194(2) (1996) 211–216.

[60] N. Van Damme, D. Baeten, M. De Vos, P. Demetter, D. Elewaut, H. Mielants, G. Verbruggen, C. Cuvelier, E.M. Veys, F. De Keyser, Chemical agents and enzymes used for the extraction of gut lymphocytes influence flow cytometric detection of T cell surface markers, Journal of Immunological Methods 236(1-2) (2000) 27–35.

[61] Y.Y. Jang, S.J. Sharkis, A low level of reactive oxygen species selects for primitive hematopoietic stem cells that may reside in the low-oxygenic niche, Blood 110(8) (2007) 3056–63.

[62] K. Naka, T. Muraguchi, T. Hoshii, A. Hirao, Regulation of reactive oxygen species and genomic stability in hematopoietic stem cells, Antioxid Redox Signal 10(11) (2008) 1883–94.

[63] A. Ludin, S. Gur-Cohen, K. Golan, K.B. Kaufmann, T. Itkin, C. Medaglia, X.J. Lu, G. Ledergor, O. Kollet,T. Lapidot, Reactive oxygen species regulate hematopoietic stem cell self-renewal, migration and development, Tas well as their bone marrow microenvironment, Antioxid Redox Signal 21(11) (2014) 1605–19.

[64] K. Ito, A. Hirao, F. Arai, K. Takubo, S. Matsuoka, K. Miyamoto, M. Ohmura, K. Naka, K. Hosokawa, Y. Ikeda, T. Suda, Reactive oxygen species act through p38 MAPK to limit the lifespan of hematopoietic stem cells, Nat Med 12(4) (2006) 446–51.

[65] G.A. Challen, N. Boles, K.K. Lin, M.A. Goodell, Mouse hematopoietic stem cell identification and analysis, Cytometry A 75(1) (2009) 14–24.

[66] G.J. Spangrude, S. Heimfeld, I.L. Weissman, Purification and characterization of mouse hematopoietic stem cells, Science 241(4861) (1988) 58–62.

[67] T.H. Hsu, S.J. Chiang, Y.H. Chu, TQuartz Crystal Microbalance Analysis of Diels-Alder Reactions of Alkene Gases to Functional Ionic Liquids on Chips, Anal Chem 88(22) (2016) 10837–10841.

[68] J. Valdez, C.D. Cook, C.C. Ahrens, A.J. Wang, A. Brown, M. Kumar, L. Stockdale, D. Rothenberg, K. Renggli, E. Gordon, D. Lauffenburger, F. White, L. Griffith, On-demand dissolution of modular, synthetic extracellular matrix reveals local epithelial-stromal communication networks, Biomaterials 130 (2017) 90–103.

[69] M.T. Ngo, B.A. Harley, The Influence of Hyaluronic Acid and Glioblastoma Cell Coculture on the Formation of Endothelial Cell Networks in Gelatin Hydrogels, Adv Healthc Mater 6(22) (2017).

[70] S. Pedron, E. Becka, B.A. Harley, Regulation of glioma cell phenotype in 3D matrices by hyaluronic acid, Biomaterials 34(30) (2013) 7408–17.

[71] S.G. Zambuto, J.F. Serrano, A.C. Vilbert, Y. Lu, B.A.C. Harley, S. Pedron, Response of neuroglia to hypoxia-induced oxidative stress using enzymatically crosslinked hydrogels, MRS Commun 10(1) (2020) 83–90.

[72] S. Okada, H. Nakauchi, K. Nagayoshi, S.I. Nishikawa, Y. Miura, T. Suda, Invivo and Invitro Stem-Cell Function of C-Kit-Positive and Sca-1-Positive Murine Hematopoietic-Cells, Blood 80(12) (1992) 3044–3050.

[73] C. McQuin, A. Goodman, V. Chernyshev, L. Kamentsky, B.A. Cimini, K.W. Karhohs, M. Doan, L. Ding, S.M. Rafelski, D. Thirstrup, W. Wiegraebe, S. Singh, T. Becker, J.C. Caicedo, A.E. Carpenter, CellProfiler 3.0: Next-generation image processing for biology, PLoS Biol 16(7) (2018) e2005970.

[74] S.S. Shapiro, M.B. Wilk, An analysis of variance test for normality (complete samples), Biometrika 52(3-4) (1965) 591–611.

[75] A. Ghasemi, S. Zahediasl, Normality tests for statistical analysis: a guide for non-statisticians, Int J Endocrinol Metab 10(2) (2012) 486–9.

[76] M.B. Brown, A.B. Forsythe, Robust Tests for the Equality of Variances, Journal of the American Statistical Association 69(346) (1974) 364–367.

[77] R.J. Carroll, H. Schneider, A note on levene’s tests for equality of variances, Statistics & Probability Letters 3(4) (1985) 191–194.

[78] R. Team, RStudio: Integrated Development Environment for R, RStudio, Inc., 2020.

[79] G.T. Hermanson, Chapter 3 - The Reactions of Bioconjugation, in: G.T. Hermanson (Ed.), Bioconjugate Techniques (Third Edition), Academic Press, Boston, 2013, pp. 229–258.

[80] B.H. Lee, H. Shirahama, N.-J. Cho, L.P. Tan, Efficient and controllable synthesis of highly substituted gelatin methacrylamide for mechanically stiff hydrogels, RSC Advances 5(128) (2015) 106094–106097.

[81] C. Claassen, M.H. Claassen, V. Truffault, L. Sewald, G.E.M. Tovar, K. Borchers, A. Southan, Quantification of Substitution of Gelatin Methacryloyl: Best Practice and Current Pitfalls, Biomacromolecules 19(1) (2018) 42–52.

[82] I. Fleming, D. Williams, Spectroscopic Methods in Organic Chemistry, 2019.

[83] E. Axpe, D. Chan, G.S. Offeddu, Y. Chang, D. Merida, H.L. Hernandez, E.A. Appel, A Multiscale Model for Solute Diffusion in Hydrogels, Macromolecules 52(18) (2019) 6889–6897.

[84] C. Mantel, S. Messina-Graham, A. Moh, S. Cooper, G. Hangoc, X.-Y. Fu, H.E. Broxmeyer, Mouse hematopoietic cell–targeted STAT3 deletion: stem/progenitor cell defects, mitochondrial dysfunction, ROS overproduction, and a rapid aging–like phenotype, Blood 120(13) (2012) 2589–2599.

[85] L. Shao, H. Li, S.K. Pazhanisamy, A. Meng, Y. Wang, D. Zhou, Reactive oxygen species and hematopoietic stem cell senescence, International Journal of Hematology 94(1) (2011) 24–32.

[86] A.K. Nguyen, P.L. Goering, V. Reipa, R.J. Narayan, Toxicity and photosensitizing assessment of gelatin methacryloyl-based hydrogels photoinitiated with lithium phenyl-2,4,6- trimethylbenzoylphosphinate in human primary renal proximal tubule epithelial cells, Biointerphases 14(2) (2019) 021007.

[87] D. Pagoria, A. Lee, W. Geurtsen, The effect of camphorquinone (CQ) and CQ-related photosensitizers on the generation of reactive oxygen species and the production of oxidative DNA damage, Biomaterials 26(19) (2005) 4091–9.

[88] J.E. Eastoe, The amino acid composition of mammalian collagen and gelatin, Biochem J 61(4) (1955) 589–600.

[89] J.D. Weaver, D.M. Headen, M.D. Hunckler, M.M. Coronel, C.L. Stabler, A.J. Garcia, Design of a vascularized synthetic poly(ethylene glycol) macroencapsulation device for islet transplantation, Biomaterials 172 (2018) 54–65.

[90] B.H. Hu, J. Su, P.B. Messersmith, Hydrogels cross-linked by native chemical ligation, Biomacromolecules 10(8) (2009) 2194–200.

[91] J. Zhu, Bioactive modification of poly(ethylene glycol) hydrogels for tissue engineering, Biomaterials 31(17) (2010) 4639–56.

[92] L.E. Jansen, N.P. Birch, J.D. Schiffman, A.J. Crosby, S.R. Peyton, Mechanics of intact bone marrow, J Mech Behav Biomed Mater 50 (2015) 299–307.

[93] E. Cambria, K. Renggli, C.C. Ahrens, C.D. Cook, C. Kroll, A.T. Krueger, B. Imperiali, L.G. Griffith, Covalent Modification of Synthetic Hydrogels with Bioactive Proteins via Sortase-Mediated Ligation, Biomacromolecules 16(8) (2015) 2316–26.

[94] H. Mao, S.A. Hart, A. Schink, B.A. Pollok, Sortase-mediated protein ligation: a new method for protein engineering, J Am Chem Soc 126(9) (2004) 2670–1.

[95] M.W. Popp, H.L. Ploegh, Making and breaking peptide bonds: protein engineering using sortase, Angew Chem Int Ed Engl 50(22) (2011) 5024–32.

[96] S. Tsukiji, T. Nagamune, Sortase-mediated ligation: a gift from Gram-positive bacteria to protein engineering, Chembiochem 10(5) (2009) 787–98.

[97] Worthington Biochemical Online Tissue Dissociation Guide, 2011. http://www.worthington-biochem.com/tissuedissociation/basic.html.

[98] W.P. Daley, S.B. Peters, M. Larsen, Extracellular matrix dynamics in development and regenerative medicine, J Cell Sci 121(Pt 3) (2008) 255–64.

[99] G.S. Schultz, J.M. Davidson, R.S. Kirsner, P. Bornstein, I.M. Herman, Dynamic reciprocity in the wound microenvironment, Wound Repair Regen 19(2) (2011) 134–48.

